# Host-microbe-immune interactions in an air-liquid interface airway model

**DOI:** 10.64898/2026.01.15.699662

**Authors:** Alexander F. Melanson, Annika Hettich, Claudia Antonella Colque, Jenny J. Persson, Pablo Laborda, Signe Lolle, Søren Molin, Helle Krogh Johansen

## Abstract

**Background:** Air-liquid interface (ALI) cell culture systems have improved the study of host-microbe interactions in respiratory infections. However, most ALI models lack immune components, limiting their ability to capture epithelial-immune crosstalk. To address this, we developed a dual-cell ALI model incorporating human peripheral blood monocyte-derived macrophages beneath differentiated airway epithelial cells.

**Methodology:** Macrophages were seeded on the basolateral side of transwell inserts using fibronectin coating. Model characterization included transepithelial electrical resistance (TEER) to assess epithelial barrier integrity, IL-8 secretion as a marker of epithelial inflammatory signaling, and confocal microscopy to evaluate cellular architecture before and after infection. Mono-and dual-cell cultures were infected with the laboratory strain *Pseudomonas aeruginosa* PAO1.

**Results:** Macrophages adhered stably to the basolateral surface without compromising epithelial barrier integrity. Following infection, IL-8 secretion was elevated in epithelial monocultures compared to dual-cell cultures, suggesting early immune modulation in the presence of macrophages. While overall bacterial burden was comparable, confocal imaging revealed clustered bacterial growth in monocultures and a more dispersed spatial distribution in dual-cell cultures.

**Conclusions:** This dual-cell ALI model enables investigation of early epithelial-immune interactions, inflammatory modulation, and bacterial colonization dynamics during airway infection. The system provides a versatile and human-relevant platform for studying respiratory host-pathogen interactions.

## Introduction

The rise of antibiotic-resistant bacteria has made persistent infections an increasingly serious threat to human health (Kavanagh, 2025; Mestrovic *et al*., 2025). There is therefore an urgent need to improve current infection models to accurately replicate the infected tissue environment for a more physiologically relevant understanding of the processes and mechanisms governing bacterial persistence. Traditional 2D cell culture systems have allowed the study and cultivation of individual human cell types, providing insights into the specific cellular responses to bacterial infection. Over the past 10-15 years, 3D cell culture model systems have been developed to bridge the gap between 2D culture and animal models (Baldassi, Gabold and Merkel, 2021a; McCarron, Parsons and Donnelley, 2021; Bauer *et al*., 2022; Moro *et al*., 2024). These systems enable researchers to better replicate the complex environments found in human tissues, thereby allowing studies of infections that closer align with physiological conditions. In the context of human airways, one commonly used model is the air-liquid interface (ALI) system, which utilizes a cell culture insert in conjunction with traditional culture dishes. These inserts provide an elevated culture surface, which supports the differentiation of airway epithelial cells into a pseudostratified epithelium, comprising basal, goblet, and ciliated cells (Colque *et al*., 2025). Moreover, the ALI model recapitulates key lung features, including mucus secretion, ciliary movement, and tight junction formation. In addition, primary cells can be derived from patient samples, allowing for a more personalized investigation strategy.

Previous studies using the ALI model to investigate infections with *Pseudomonas aeruginosa, Staphylococcus aureus,* and SARS-CoV-2 have examined immune responses, microbial colonization, epithelial barrier integrity, and molecular mechanisms underlying host-pathogen interactions (Baldassi, Gabold and Merkel, 2021b; Ciszek-Lenda *et al*., 2023; La Rosa, Molin and Johansen, 2025).

However, traditional ALI models usually lack immune cell components, limiting their utility to model immune interactions during infection. Given that immune cells are both resident and actively recruited, and bacteria need to evade their action to establish an infection, their inclusion is critical for obtaining a more realistic picture of host-pathogen interactions during infection. Macrophages were chosen for this model due to their pivotal role in the innate immune response as they are important pathogen-recognizing phagocytes constitutively present in tissues, and because they participate in both acute and chronic infections (Mitchell*, Chen* and Portnoy, 2016; Macpherson *et al*., 2017; Acosta and Alonzo, 2023; Ciszek-Lenda *et al*., 2023; Peña-Cearra *et al*., 2024; Sit *et al*., 2024). In mono-culture studies, macrophages infected with pathogens such as *Mycobacterium tuberculosis*, *Salmonella enterica*, and *P. aeruginosa,* have demonstrated altered intracellular signaling and spatial dynamics of bacterial colonization (Nix *et al*., 2007; Fallows *et al*., 2016; Maphasa, Meyer and Dube, 2020; Moser *et al*., 2021; Ciszek-Lenda *et al*., 2023; Nickerson *et al*., 2024). Despite this, the complex interactions between airway epithelial cells, macrophages, and bacteria remain poorly understood.

To address this, we developed a dual-cell ALI model that incorporates human peripheral blood monocyte-derived macrophages on the basolateral side of a differentiated airway epithelial layer. Macrophages were seeded to the basolateral side of cell culture inserts to mimic the homeostatic environment before infection. This model was infected with *P. aeruginosa*, a prevalent opportunistic pathogen able to cause persistent infections specially in the airways of people with structural lung diseases(Wood, Kuzel and Shafikhani, 2023). This study aimed to (1) characterize macrophage adhesion to the cell culture insert, (2) assess the impact of macrophage presence on epithelial integrity, (3) quantify immune signaling via IL-8 secretion, and (4) investigate *P. aeruginosa* colonization, growth, and spatial organization in the presence and absence of macrophages.

## Materials and Methods

### Preparation of cell culture inserts, BCi-NS1.1 cell line, and *Pseudomonas aeruginosa* PAO1

For the cell culture system, we used 1 μm pore polyester membrane inserts (Falcon). BCi-NS1.1 cell line was expanded and seeded into transwells and matured for 28 days as previously described (Walters *et al*., 2013; Laborda *et al*., 2024). To prepare the culture inserts for proper cell adhesion, type I collagen (Gibco) was added to the apical side of the inserts and incubated for 30 min at room temperature (RT). Subsequently, inserts were inverted and 20µL fibronectin (Merck) at 5 µg/mL was added to the basolateral side and incubated for 2 h at 37 °C. For fibronectin coating optimization, two commonly used diluents were tested: (1) Dulbecco’s Modified Eagle Medium (DMEM, Gibco) and (2) Phosphate buffered saline (PBS, Gibco).

Preparation of the laboratory strain *P. aeruginosa* PAO1 inoculum and the infection procedure in the ALI model followed the protocol described by Laborda et al (Laborda *et al*., 2024). Two minor modifications were made: (1) the infecting inoculum was reduced from 1000 to 100 colony forming units (CFU), and (2) the infection duration was extended from 14 to 16 h.

### Preparation of macrophages

Peripheral blood mononuclear cells (PBMCs) were isolated from fresh buffy coats obtained from healthy donors through the Department of Clinical Immunology, the Blood Bank, at Rigshospitalet, Copenhagen, Denmark. Buffy coats from 3-4 donors were pooled for each experiment to increase cell yield and minimize the impact of donor-specific variability. PBMCs were processed on the same day of collection. In brief, 50 mL of buffy coat was diluted 1:1 with PBS supplemented with 2 % fetal-bovine-serum (FBS, Gibco). PBMCs were isolated using 50 mL LEUCOSEP separation tubes (GREINER) pre-filled with Lymphoprep (STEMCELL Technologies). Following isolation, cells were washed twice in PBS + 2 % FBS with 2mM Ethylenediaminetetraacetic acid (EDTA) and centrifuged at 120 x g for 10 min at 4 °C to remove residual platelets. The resulting PBMCs were cryopreserved at a concentration of 1 x 10^7^ cells/mL in FBS supplemented with 10 % dimethyl-sulfoxide (DMSO, Thermo Fisher Scientific).

Cryopreserved PBMCs were quickly thawed in a 37 °C water bath and immediately transferred into pre-warmed M0 base media consisting of RPMI 1640 (Gibco) supplemented with 10 % FBS and 50 ng/mL macrophage colony-stimulating factor (M-CSF, Miltenyi Biotec). Monocytes were purified via negative selection using the MojoSort™ Human Pan Monocyte Isolation Kit (Biolegend), following the manufacturer’s instructions. Purified monocytes were seeded at a density of 14 x 10^6^ cells/mL in M0 base media in P-100 tissue culture-treated dishes and incubated at 37 °C with 5 % CO_2_. Cells were differentiated into macrophages over 6 days, with a media change at 24 h to remove non-adherent cells.

For macrophage harvesting, TrypLE (Gibco) was applied to the culture plates and incubated at 37 °C for 5-10 min. Cells were lifted using a cell scraper and cold PBS was added to inactivate TrypLE and prevent macrophage adherence to collection tubes. Finally, cells were pelleted by centrifugation at 300 x g for 5-10 min and resuspended to the desired concentration for downstream processing and analysis.

### Establishing the dual-cell culture model

Airway epithelial cells, fully differentiated on the apical side of a transwell, were used to establish the dual-cell ALI model. Prior to macrophages seeding, TEER was measured to confirm epithelial integrity and media from the basolateral compartment was removed. To enable macrophage adhesion, culture inserts were inverted, and for the initial optimization fibronectin (Merck) was diluted in either DMEM or PBS. On day 0, fibronectin (5 µg/mL) was added to the basolateral side of the inserts and incubated for 2 h at 37 °C with 5 % CO_2_. On day 27, fibronectin (5 µg/mL) was reapplied, and incubated for 30 min. Following this, 20 µL of macrophage suspension (containing 8.5 x 10^4^ cells), was applied to the fibronectin-coated basolateral side and incubated for 1 h at 37 °C with 5 % CO_2_. Control inserts without macrophages were treated identically, with macrophage-free media added to the basolateral side.

After incubation, media was aspirated and the inserts were carefully returned to their original orientation in the same 24-well plates. Each well contained 400 µL of pre-warmed media composed of a 1:1 mixture of M0 base media and Pneumacult-ALI maintenance media (STEMCELL Technologies), supplemented with 4 µg/mL heparin, 480 ng/mL hydrocortisone, Pneumacult-ALI 10x supplement, and Pneumacult-ALI maintenance supplement (all from STEMCELL Technologies). In control wells lacking epithelial cells, 50 µL of 1:1 mixture media was added apically to maintain comparable conditions. All cultures were incubated at 37 °C in a humidified 5 % CO_2_ incubator until the time of infection.

### Immunostaining and confocal microscopy of dual-cell culture model

Before staining, both the apical and basolateral compartments of the transwells were rinsed once with PBS. Cells were then fixed by adding 4 % paraformaldehyde (PFA, Thermo Fisher Scientific) to the apical (200 µL) and basolateral (400 µL) compartments and incubated for 20 min at 4 °C. After fixation, PFA was removed, and samples were washed twice with PBS. Fixed membranes were excised from the culture inserts with a surgical scalpel and placed on a parafilm-covered rectangular petri dish. Membranes were then blocked with 100 µL of 3 % goat serum (Jackson Immuno) in PBS for 1 h at RT. Droplet-based staining ensured that both the apical and the basolateral surfaces remained in contact with solution, thereby enhancing visualization of macrophages attached to the basolateral side. Primary antibodies were diluted in blocking buffer and applied to membranes in 100 µL droplets. Membranes were incubated overnight at 4 °C with rabbit anti-CD14 (Thermo Fisher Scientific) and acetylated alpha-tubulin (Santacruz Biotechnology) antibodies. Following incubation, membranes were washed three times with PBS. Secondary staining was performed using goat anti-rabbit antibody (Thermo Fisher Scientific) diluted in blocking buffer, applied in 100 µL droplets, and incubated for 1 h at RT. For intracellular staining, cells were permeabilized using 1 % saponin (Sigma Aldrich) in blocking buffer applied as 100 µL droplets for 30 min at RT. Nuclei were counterstained with DAPI for 10 min at RT, followed by three PBS washes, prior to antibody application.

For mounting, membranes were placed on glass slides, covered with a drop of VECTASHIELD® Antifade Mounting media (VWR), and sealed with a coverslip using nail polish. Confocal images were acquired using a Leica Stellaris 8 confocal microscope (40x oil immersion objective, NA 1.3).

### Flow cytometry and staining of airway epithelial cells

Basal cell differentiation was assessed after 28 days of ALI culture. To remove accumulated mucus, the apical side of each transwell was treated with 200 µL of PBS containing 10 mM dithiothreitol (DTT, Sigma Aldrich) and incubated for 10 min at 37 °C in a 5 % CO_2_ humidified incubator. Residual DTT was removed by three washes with 75 µL of PBS. The media from the basolateral compartment was aspirated and enzymatic dissociation of the cell layer was performed using trypsin (Gibco) supplemented with 0,5 mM EDTA. A volume of 500 µL was added to the basolateral compartment and 200 µL to the apical compartment. Cells were incubated for 10-20 min at 37 °C in a 5 % CO_2_ humidified incubator. Apical compartment cells were harvested by pipetting and transferred to a 1.5 mL Eppendorf tube. The basolateral trypsin solution was then used to rinse the insert twice to maximize cell recovery. To stop trypsin activity, 500 µL of complete media (RPMI 1640 with 20 % FBS) was added to each tube. The cell suspensions were filtered through a 100 µm filter to reduce clumping and collected in a 15 mL Falcon tube containing 3 mL of complete media. The filter was then rinsed with 1 mL of complete media to ensure maximal cell recovery. Cells were centrifuged at 400 x g for 5 min at RT. The supernatant was removed, and the cell pellet resuspended in 1 mL of 0.5 % (v/v) PFA diluted in PBS, transferred to a clean 1.5 mL Eppendorf tube, and incubated for 8 min at RT. Cells were subsequently centrifuged at 300 x g for 5 min and washed twice with PBS to remove residual PFA. Fixed cells were either used immediately for flow cytometry analysis or stored at-80 °C for later use.

For staining, cells were resuspended in 100 µL staining buffer consisting of 5 % (v/v) goat serum (Jackson Immuno) in PBS and incubated for 15 min at RT on a rotator (350 x g) to block non-specific primary antibody binding, including Fc receptor interactions. Antibodies against the surface proteins CD271 and CD66c (Thermo Fischer Scientific) were added directly to the blocked cells and incubated for 30 min at RT on a rotator (350 x g), protected from light. After surface staining, cells were washed three times with 700 µL of PBS. For intracellular staining, cells were permeabilized in a 100 µL staining buffer containing 0.2 % (w/v) saponin (Sigma Aldrich) for 15 min at RT on a rotator (350 x g). Antibodies targeting intracellular markers Muc5AC and TUBA (Thermo Fisher Scientific) were then added directly to the permeabilized cells and incubated for 30 min at RT on a rotator (350 x g), protected from light.

After staining, cells were washed three times in 700 µL PBS supplemented with 0.1 % saponin and resuspended in 250 µL PBS containing 0.1 % bovine serum albumin (Sigma-Aldrich). The stained cells were transferred to a flat-bottom 96-well plate (Greiner). Samples were analyzed using an Agilent Novocyte flow cytometer. To correct for fluorescence spillover, spectral overlap compensation was performed using the Mouse Ig, κ/Negative Control Compensation Particles Set (BD Biosciences), with each antibody used in the experiment. Unstained cells were included as reference.

### Flow cytometry analysis of macrophages

Harvested macrophages were stained with fixable viability dye eFluor 450 (Thermo Fisher Scientific), followed by fixation in 4.2 % PFA (BD Bioscience), for 15 minutes at RT. Non-specific Fc receptor interactions were blocked using human Fc blocking reagent (BD Bioscience) for 30 min at RT, and cells were stained with anti-CD16-PE and anti-CD14-FITC (Thermo Fisher Scientific). Samples were analzyed using a NovoCyte Quanteon flow cytometer (Agilent) using ultracomp ebeads^tm^ (Thermo Fisher Scientific) for compensations, as well as fluorescent minus one (FMO) were also used as a compensation control.

### Quantification of infection parameters after *P. aeruginosa* infection

Bacterial load (CFU/mL) in the apical, attached, and basolateral compartments of ALI transwells was measured by plating serial fold dilutions of each sample in triplicates in LB agar plates. In addition, TEER and IL-8 secretion after infection with *P. aeruginosa* were measured. All methods listed above were conducted as previously described (Laborda et al., 2024).

## Data analysis

Flow cytometry data was analyzed using the Flow Cytometry Analysis Software Flowlogic (Version 7.3). Confocal images were analyzed and edited using the Fiji (software based on Windows Image J). Z-stack images obtained by confocal microscopy were Z-projected using the maximum intensity projection. All plots and statistics were generated with GraphPad Prism Version 10.5.0 for Mac (GraphPad Software, San Diego, California USA). Data is presented as mean ± SEM from three separate biological replicates, each with three technical replicates. Statistical significance was determined by t-test or one-way ANOVA with Tukey’s multiple comparisons test and is indicated as *p ≤ 0.05, **p ≤ 0.01, ***p ≤ 0.001 and ****p < 0.0001.

## Results

### Macrophage adhesion to cell culture inserts

The primary objective of this study was to establish a dual-cell culture model incorporating monocyte-derived macrophages to an airway epithelial cell culture system. To achieve this, macrophage adhesion to culture inserts was first evaluated. Human monocytes were differentiated into macrophages over six days in the presence of macrophage colony-stimulating factor (M-CSF). Brightfield microscopy confirmed the expected elongated morphology of adherent macrophages (Figure 1A). Surface marker analysis by flow cytometry showed robust expression of CD14 and CD16, with similar patterns of marker expression as previously described *(Buckner* et al.*, 2011; Lee* et al.*, 2020)*, confirming successful macrophage differentiation (Figure 1B).

**Figure 1.**
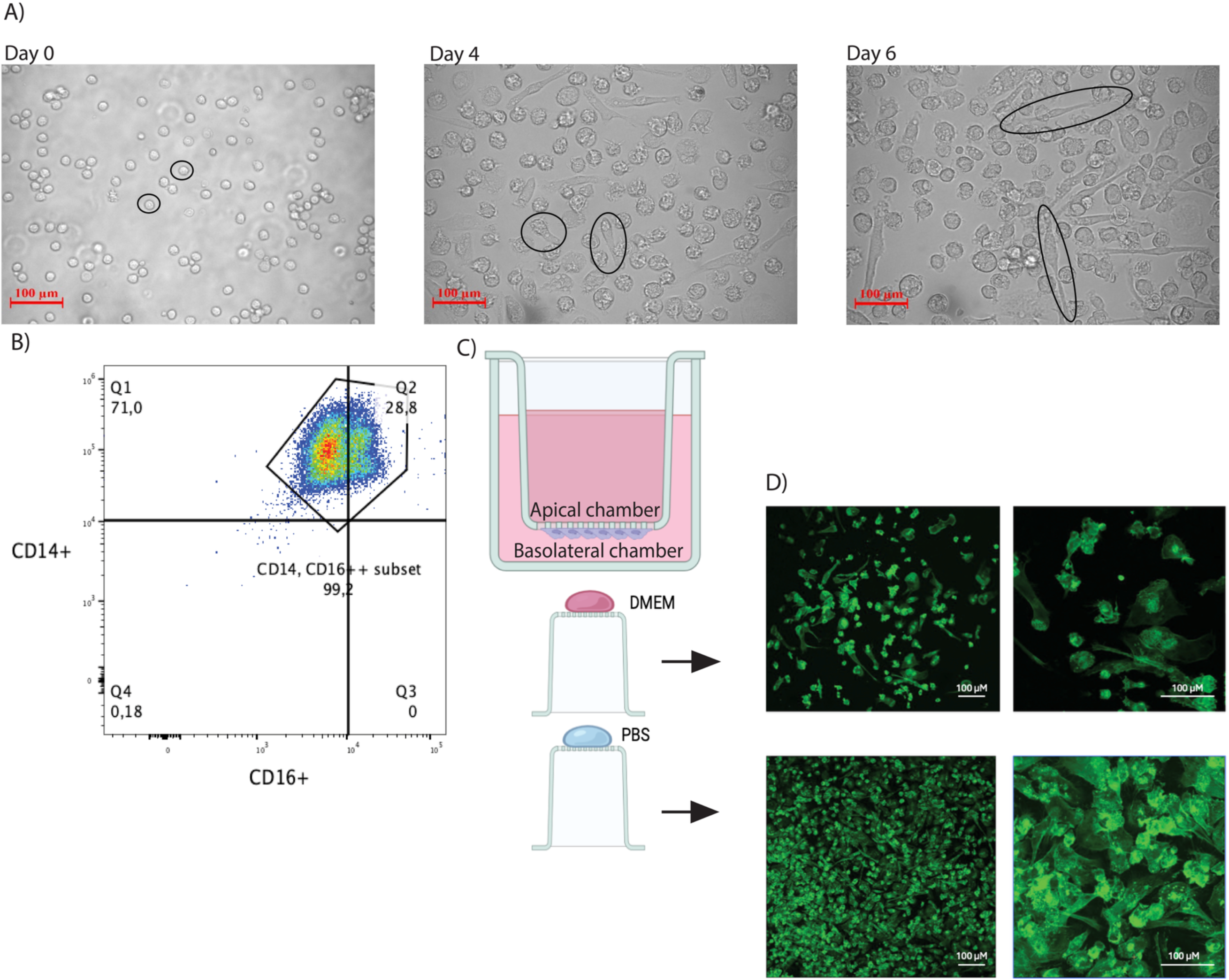
**Maturation and phenotype development of human peripheral monocyte-derived macrophages and adhesion to cell culture inserts**. A) Development of monocytes into macrophage-like cells during *in vitro* culture. Light microscopy images (magnification: 20x) of cellular phenotypes at day 0 (after negative selection), day 4, and day 6. B) Flow cytometry-based characterization of macrophages. C) Schematic cartoon of macrophage seeding to the basolateral side of the cell culture inserts, image was created in BioRender *by Alexander F. Melanson* (2025) https://BioRender.com/lwgs8nj. D) Adherence of macrophages to cell culture inserts coated with fibronectin diluted in Dublecco’s Modified Eagle Medium (DMEM, top) or Phosphate Buffered Saline (PBS, bottom). Confocal microscopy images of macrophages stained with anti-CD14 (green). Data are representative of three biological replicates, each with three technical replicates.

Macrophages were seeded onto the basolateral side of cell culture inserts (Figure 1C) coated with human plasma fibronectin diluted in either DMEM or PBS to identify the most efficient fibronectin coating diluent (Figure 1D). Confocal microscopy of CD14-stained macrophages showed cellular adhesion in both conditions, although visual assessment suggested fibronectin diluted in PBS better supported this feature (Figure 1D).

### Establishment of dual-cell culture model

The sequential steps from ALI model differentiation, macrophage adhesion, to dual-cell culture establishment is displayed in Figure 2A. In brief, human airway basal cells were seeded and differentiated on cell culture inserts. After 21 days of ALI differentiation, human monocyte-derived macrophages were generated by negatively selected monocytes, then differentiating them 6 days in M0 base media. On day 27, one day prior to starting the infection, macrophages were seeded to the basolateral side of the cell culture inserts.

**Figure 2.**
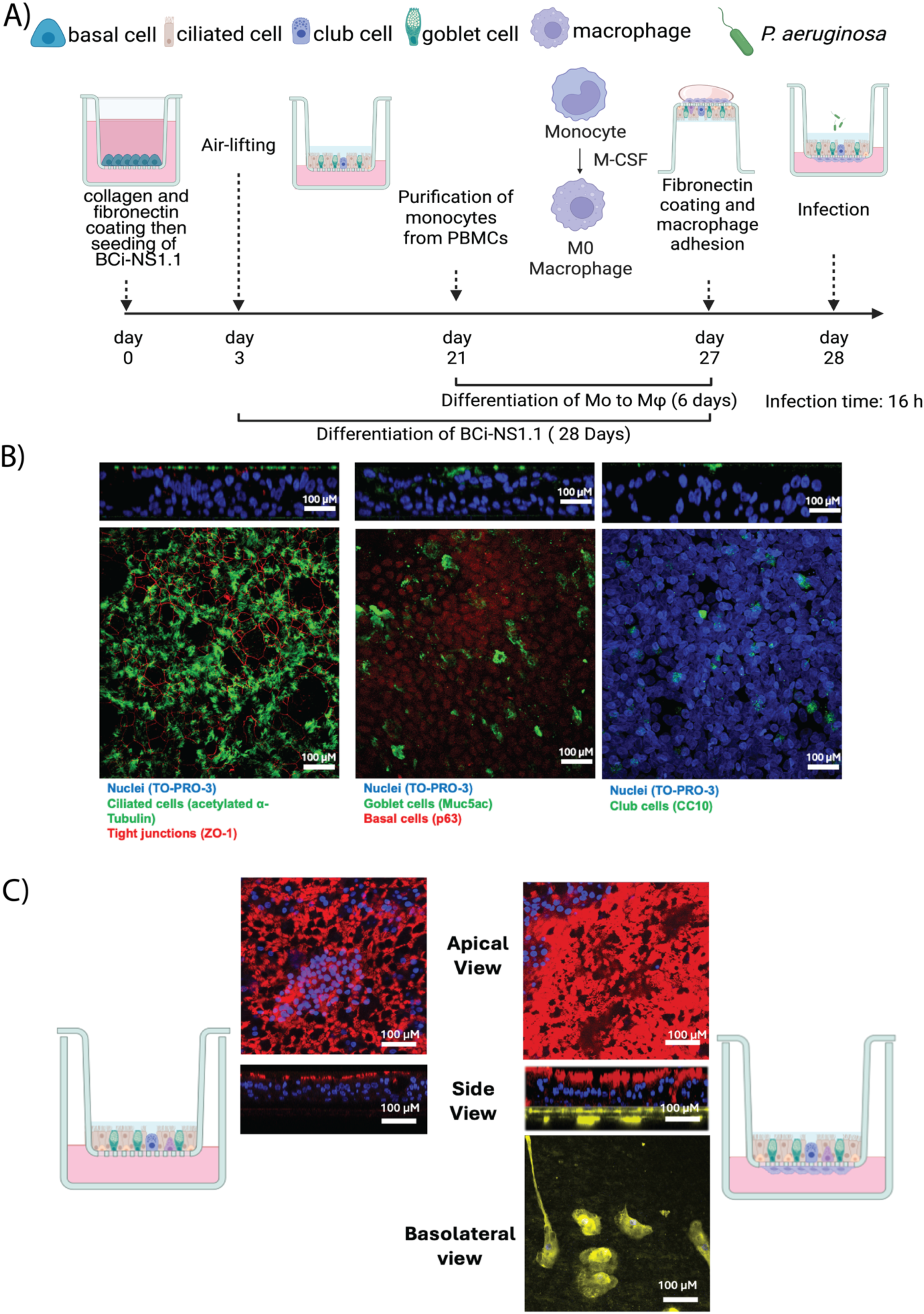
Establishment of the dual-cell culture infection model. A) Schematic timeline of the process to establish the model image was created in BioRender by Alexander F. Melanson (2025) https://BioRender.com/lwgs8nj B) Confocal images of BCi-NS1.1 cells: left panel shows staining of cell nuclei (blue, To-pro-3), ciliated cells (green, acetylated alpha-tubulin), and tight junctions (red, ZO-1); middle image, nuclei (blue, To-pro-3), goblet cells (green, Muc5ac), and basal cells (red, p63); right panel, nuclei (blue, To-pro-3) and club cells (green, CC10). C) Left, first a schematic cartoon of a cell culture insert system without macrophages, then a mono-cell culture insert with only BCi-NS1.1 cells with ciliated cells (red, acetylated alpha-tubulin), and nuclei (blue, DAPI). Right, dual-cell culture model, with macrophages (yellow, CD14) adhering to the basolateral side of the cell culture insert. Then another schematic cartoon, this one showing macrophages adhered to the basolateral side of transwells. Apical view for both left and right images are a maximum projection of the Z-stacks.

To assess the cellular composition of the epithelial layer, transwell inserts containing fully differentiated airway epithelial BCi-NS1.1 cells were stained and analyzed by confocal microscopy (Figure 2B). Differentiation led to a fully mature pseudostratified epithelial tissue, characterized by the formation of tight junctions and the presence of multiple cell types representative of an airway epithelium (Supplementary Figures 1 A and 1 B).

Figure 2C shows the mono-culture model (epithelial tissue only) and the dual-cell culture model (epithelial tissue and macrophages). Because of the relative instability of fibronectin, the wells were supplemented with fresh fibronectin prior to macrophage seeding to ensure their adhesion, which was confirmed by confocal imaging (Figure 2D). To evaluate the retention of adherent cells, detached macrophages were collected and counted 24 h after seeding them. No significant loss of macrophages was observed (approximately 0.006% to 0,009% macrophages detached).

Importantly, the addition of macrophages had no measurable effect on epithelial barrier function, as TEER values remained unchanged (Supplementary Figure 1C). These findings confirm that macrophages can be successfully incorporated into the basolateral side of the culture insert beneath a fully differentiated airway epithelium without compromising epithelial integrity.

### Infection of mono-and dual-cell culture model

Both mono-and dual-cell culture models were infected with 100 CFU of the laboratory strain *P. aeruginosa* PAO1 and incubated for 16 h. To test differences in epithelial immune response between the two models, IL-8 secretion was measured as a marker of ability for immune cell recruitment (Figure 3A). Under mock conditions, IL-8 secretion was significantly higher in the dual-cell model compared with the epithelial monoculture (one-way ANOVA with Tukey’s multiple comparisons test, p < 0.0001). Upon PAO1 infection, IL-8 levels increased significantly in the epithelial monoculture relative to mock controls (one-way ANOVA with Tukey’s multiple comparisons test, p < 0.05). In contrast, PAO1 infection did not result in a significant increase in IL-8 secretion in the dual-cell model. Consequently, IL-8 levels in infected epithelial monocultures were significantly higher than in infected dual-cell cultures (one-way ANOVA with Tukey’s multiple comparisons test, p < 0.05).

**Figure 3.**
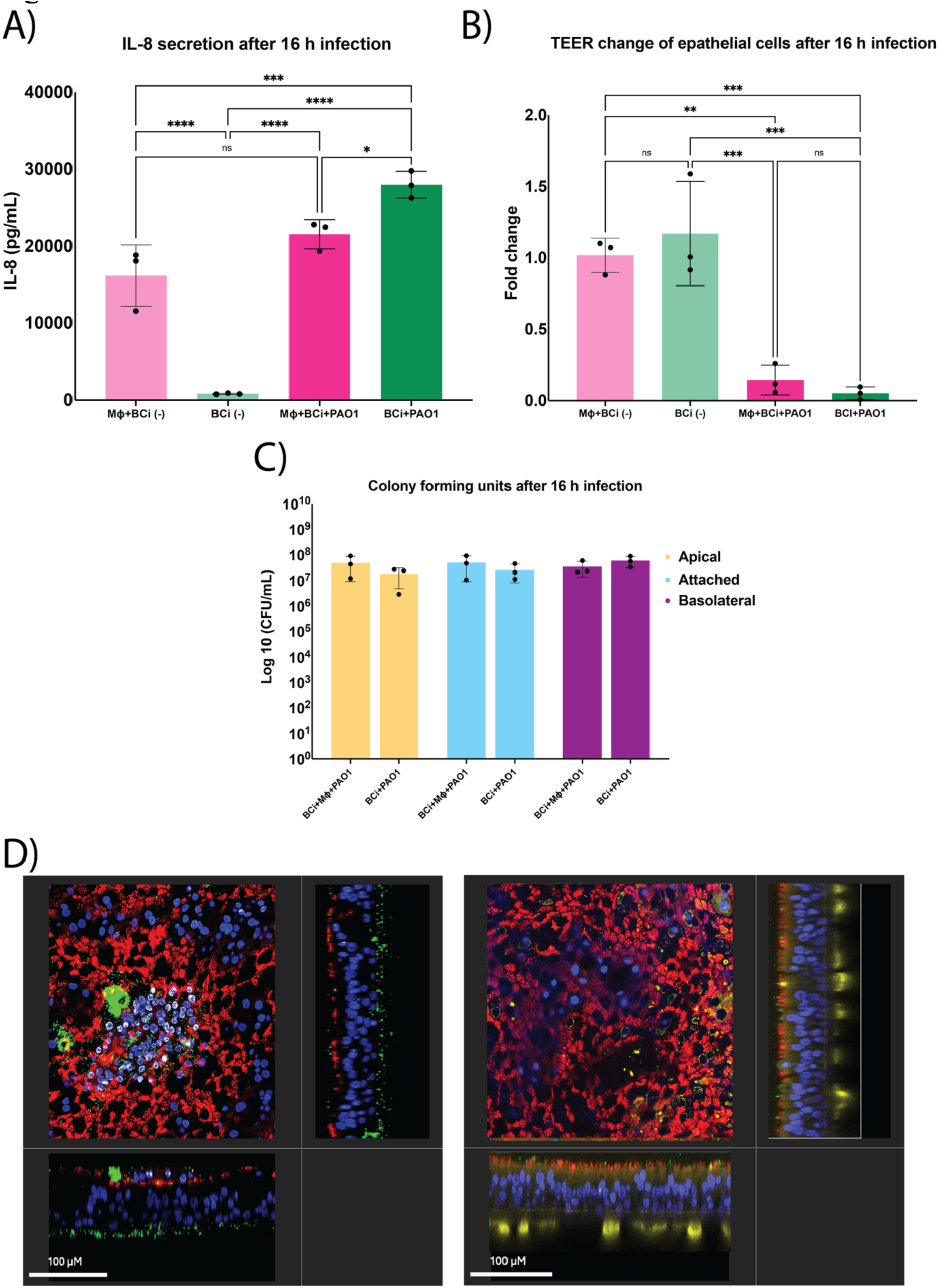
Effect of *Pseudomonas aeruginosa* infection in mono-and dual-cell culture models. A) IL-8 quantification (pg/mL) after 16 h of infection with *P. aeruginosa*. B) Fold change of Transepithelial Electrical Resistance (TEER) (Ω·cm^2^) in both models after infection with *P. aeruginosa* for 16 h. C) Bacterial load in colony forming units (CFU/mL) across the different compartments of the dual-cell ALI model, apical, attached (bacteria present within the epithelial cells or firmly attached to the surface), and the basolateral compartment.. D) Orthogonal and cross-sectional views of the mono-and dual-cell culture model after 16 h of infection with *P. aeruginosa* (scale bar: 100 µm). Left image, a mono-cell culture infection model showing ciliated cells (red, acetylated alpha-tubulin), *P. aeruginosa* (green, sfGFP), and cell nuclei (blue, DAPI); right image shows a dual-cell culture model with ciliated cells (red, acetylated alpha-tubulin), *P. aeruginosa* (green, sfGFP), cell nuclei (blue, DAPI), and macrophages (yellow, CD14). All Images were taken at 40X magnification. Data represent mean ± SEM from three biological replicates, each with three technical replicates. Statistical significance was determined by one-way ANOVA with Tukey’s multiple comparisons test for IL-8, TEER and CFU comparisons, and is indicated as *p ≤ 0.05, **p ≤ 0.001, *** p ≤ 0.0001, and **** p < 0.0001. p ≤ 0.0001, p < 0.0001

To investigate the effect of infection on the epithelial cell layer integrity, TEER was measured post-infection. (Figure 3B). Under mock conditions, no significant difference in TEER was observed between epithelial monocultures and dual-cell cultures containing macrophages.

Infection with *P. aeruginosa* PAO1 caused a significant reduction in TEER compared with corresponding mock controls in both mono-and dual-cell models (one-way ANOVA with Tukey’ s multiple comparisons test, p ≤ 0.0001). TEER values in infected epithelial monocultures were also significantly lower than those in mock dual-cell cultures (p ≤ 0.001). Importantly, no significant difference in TEER was detected between infected mono-and dual-cell models, indicating that macrophage presence does not prevent the initial epithelial barrier disruption induced by *P. aeruginosa*.

Bacterial burden in the apical, attached, and basolateral compartments, revealed comparable bacterial loads across models (Figure 3C, one-way ANOVA with Tukey’s multiple comparisons test, p>0.05). Visualization of the cell culture inserts showed (Figure 3C) large bacterial aggregates consistent with the known aggregate-forming behavior of *P. aeruginosa* (Rossi *et al*., 2021; Laborda *et al*., 2024; Leoni Swart *et al*., 2024). In contrast, in the dual-cell culture model a more dispersed bacterial distribution with predominantly single bacteria present throughout the epithelium was observed.

## Discussion

This study introduced a dual-cell ALI airway model containing epithelial cells and monocyte-derived macrophages. The aim was to establish a more realistic airway infection model system. Previous ALI models often only relied on epithelial monocultures or limited immune additions (Gras *et al*., 2017; Baldassi, Gabold and Merkel, 2021a; Paplinska-Goryca *et al*., 2021). Here, we successfully adhered macrophages into basolateral membrane of the ALI model while preserving epithelial barrier integrity and maintaining macrophage phenotype, as indicated by typical morphology and sustained CD14 expression (Hamidzadeh *et al*., 2020; Paplinska-Goryca *et al*., 2021; Hickman *et al*., 2023). This dual-cell configuration supports the study of interactions between epithelial and immune cells and enables simultaneous analysis of microbial behavior and early host responses.

Macrophages were included because they serve as key immune sentinels in the airways. Under homeostatic conditions, monocytes enter from the bloodstream and differentiate beneath the epithelium. From this position, macrophages survey the airways by extending protrusions through tight junctions and migrate into the airway lumen when infection occurs (Meeker *et al*., 2012; Chikina *et al*., 2020; Pérez-Rodríguez *et al*., 2022; Yang *et al*., 2023). To reflect this anatomy, we seeded macrophages on the basolateral membrane building on previous studies that have positioned other myeloid-derived immune cells such as dendritic cells and macrophages directly in the apical chamber (Gras *et al*., 2017; Baldassi, Gabold and Merkel, 2021a; Paplinska-Goryca *et al*., 2021; Misiukiewicz-Stępien *et al*., 2022). As a result, our model retains the natural spatial organization of airway tissue and immune cells.

Technical optimization was essential for the dual-cell culture system, and membrane pore size was one of the key parameters. Previous work has shown that very small pores (≤0.4 µm) can restrict macrophage migration and limit transepithelial probing because they prevent cells from extending their projections across the membrane (Pérez-Rodríguez *et al*., 2022; Yang *et al*., 2023; Li *et al*., 2025). In contrast, large pores such as 4 µm can weaken epithelial barrier formation. Based on these observations, we selected a 1.0 µm pore size to balance both requirements. Together, these considerations highlight how membrane properties directly influence immune–epithelial communication in ALI-based co-culture models Macrophage adhesion also required refinement. A single fibronectin coating, applied according to standard instructions, was insufficient. After testing, we added a second short incubation period of 30 min with fibronectin immediately before seeding. This step improved macrophage attachment without removing epithelial media long enough to compromise epithelial integrity. With this adjustment, macrophages remained stably adhered throughout infection experiments.

*P. aeruginosa* organization on the epithelial surface changed noticeably in the presence of macrophages. Consistent with previous reports, epithelial monocultures supported the formation of bacterial clusters (Moser *et al*., 2021; Ciofu *et al*., 2022; Laborda *et al*., 2024; Leoni Swart *et al*., 2024). However, introducing macrophages resulted in a more dispersed distribution, suggesting that early macrophage activity alters the spatial dynamics of bacterial attachment. Although our observations were qualitative, they indicate a possible change in bacterial infection phenotype. This shift in spatial organization may represent an early macrophage-mediated containment mechanism, occurring before overt phagocytosis or bacterial clearance. Quantitative spatial analysis in future work will help determine whether macrophage presence actively disrupts microcolony initiation or merely prevents their maturation.

The reduction in IL-8 when macrophages were present suggests that macrophages shape epithelial inflammatory output during the earliest stages of infection. This interpretation aligns with work showing that macrophages can attenuate epithelial chemokine release to prevent excessive neutrophil recruitment (Bishayi *et al*., 2017; Moser *et al*., 2021; Toya *et al*., 2024). Under mock conditions, macrophage inclusion led to increased IL-8 secretion, which may reflect basal immune–epithelial communication rather than a pathological inflammatory response. Complementing this, TEER measurements confirmed that baseline epithelial barrier integrity was preserved in the dual-cell model, indicating that macrophages do not disrupt epithelial structure under homeostatic conditions. Together, these findings indicate that while macrophages modulate epithelial inflammatory signaling, they do not lead to initial epithelial barrier loss. Our results highlight that epithelial responses measured in monoculture may overestimate in vivo inflammation, and underscore the importance of incorporating immune cells to capture early host-pathogen interactions. Distinguishing whether IL-8 modulation occurs through receptor-mediated sequestration, altered transcription, or contact-dependent signaling remains an important future direction.

Our model provides several advantages over standard ALI systems. It maintains physical separation between cell types while enabling direct communication. It can also be adapted to other pathogens or immune cell types. Together, these features make it suitable for studying epithelial-immune cell interactions during bacterial, viral, and fungal infections.

However, this system has some limitations. The static transwell design does not reproduce airflow, mechanical stretch, or mucus transport and longer infection periods would also have allowed us to study in-depth biofilm maturation. The macrophages that we added to the model system were blood-derived rather than fully tissue-resident. In addition, incorporation of patient-derived cells could further have supported personalized infection research.

Despite these limitations that can be addressed in future work, this dual-cell ALI model offers a realistic and controllable platform for airway infection studies. It bridges the gap between monocultures and *in vivo* systems and provides a useful tool for investigating respiratory immune responses and evaluating targeted therapeutic strategies.

## Ethical approval and consent

This study was approved by the local ethics committee of the Capital Region of Denmark (Region Hovedstaden), registration number H-20024750.

## Authors contribution

AM, AH, JP, SM and HKJ contributed to conception and design of the study. AM and AH performed the experimental work and statistical analysis. AM, AH, CAC, PK and HKJ wrote the manuscript. All authors contributed to manuscript revision, read, and approved the submitted version.

## Acknowledgements

The authors thank the staff at the blood bank at Rigshospitalet (Copenhagen, Denmark) for preparing of the buffy coats for this study. Furthermore, the authors thank Ronald G. Cristal (Weil Cornell Medical College, New York, USA) for providing the BCi-NS1.1 cell line.

---

**Supplementary Figure 1.**
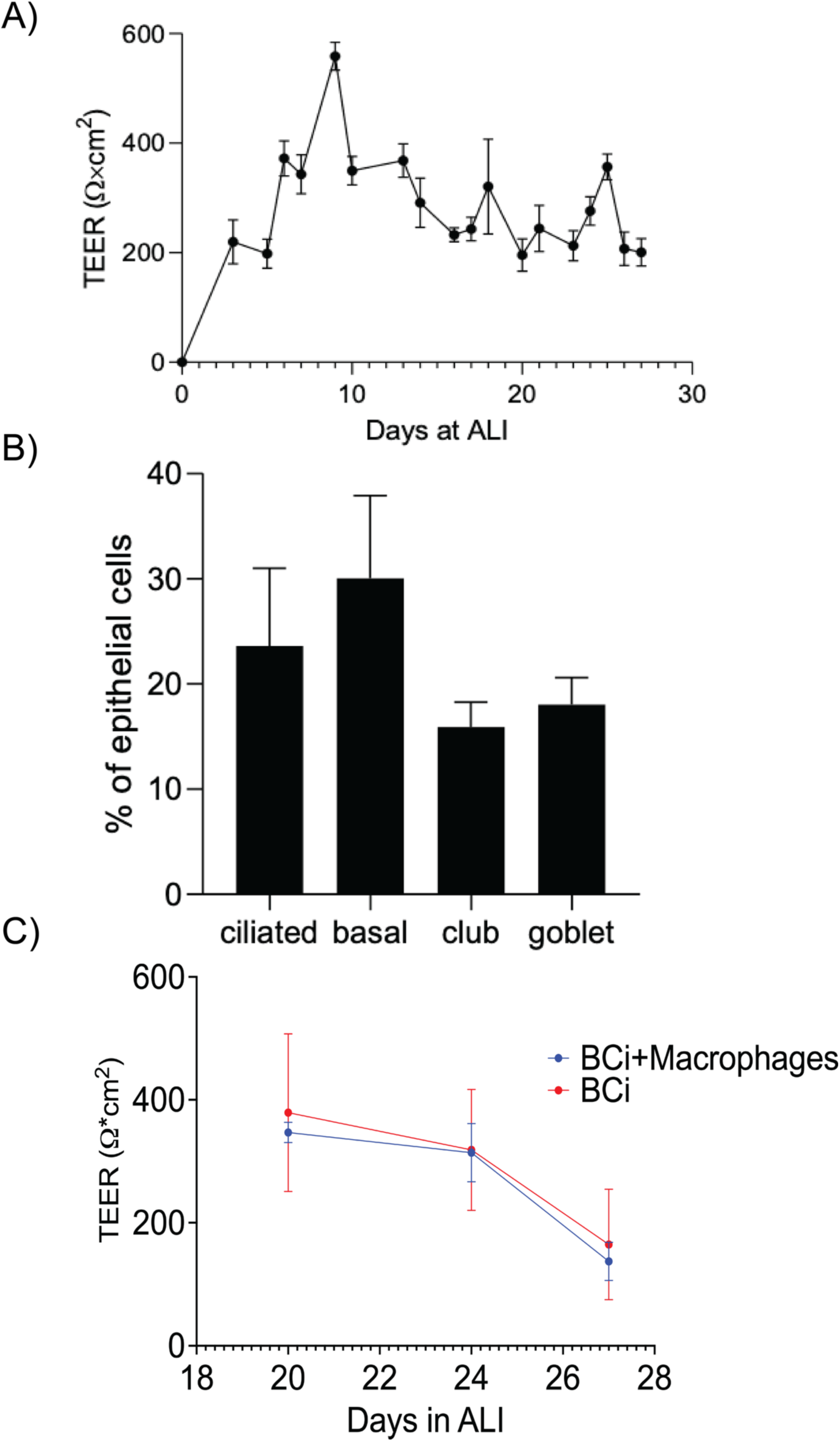
**Epithelial barrier and cell population development, and surface integrity after macrophage adhesion**. A) Changes in Trans-Epithelial-Electrical Resistance (TEER) values as the BCi-NS1.1 cells mature over 28 days. B) Percentage of cells present in the mono-cell culture model using flow cytometry. C) TEER values of mono-and dual-cell models (red: BCi-NS1.1 cells alone, blue: BCi-NS1.1+Macrophages). Data represent mean ± SEM from three biological replicates, each with three technical replicates.

